# Advancing microbial isolation: The impact of leaf mold extract agar on soil samples

**DOI:** 10.1101/2024.03.15.585253

**Authors:** Atsushi Miyashita, Kazuhiro Mikami, Masaki Ishii, Masanobu Miyauchi, Fumiaki Tabuchi

## Abstract

In this study, we developed a new agar medium using leaf mold extract and evaluated its microbial cultivation performance with soil samples. As a control for performance evaluation, the general-purpose nutrient medium YME agar was used. For YME agar, isolated microbes were confirmed to belong mainly to the *Bacillus* genus through 16S rRNA gene sequencing. In contrast, for the leaf mold agar, bacteria belonging to either *Streptomyces* or *Rhizobium* were frequently isolated. Also, of the 51 sequenced isolates on the leaf mold agar medium, an unidentified species (i.e., not listed in databases) with 16S rRNA sequence identity below 98.7% was found. The unidentified species did not grow in standard nutrient media (YME, BHI, LB10, or TSB) but grew in 10% leaf mold extract. The findings of this study suggest that the use of leaf mold agar medium can effectively isolate soil microbes that are difficult to culture in general-purpose nutrient media such as YME. Moreover, this approach indicates a viable method for discovering unknown, previously-uncultured microbial species from soil samples.

## Introduction

Throughout history, humans have successfully isolated various pharmaceuticals from the culture supernatants of soil microorganisms. These include antibiotics (e.g., streptomycin (1, 2) and micafungin (3)), antiparasitic drugs (e.g., ivermectin (4)), immunosuppressants (e.g., tacrolimus (5)), and anticancer agents (e.g., doxorubicin (6)), which have all been applied in clinical settings. More recently, lysocin E, which has shown therapeutic effects against Methicillin-resistant Staphylococcus aureus (MRSA) infections, was isolated from bacteria in soil and is being considered as a promising new antibiotic (7, 8). It is believed that among the microorganisms that have not yet been cultured, many may produce unknown compounds that could potentially serve as pharmaceutical candidates. To continue discovering valuable compounds from environmental microorganisms, we must deepen our understanding of microbial cultivation techniques.

Our surrounding environment, particularly soil, hosts a rich ecosystem composed of a vast array of microorganisms. According to research by van der Heijden et al., it is estimated that there are between 10^10^ to 10^11^ bacteria in just one gram of soil (9). This suggests that soil is particularly abundant in microbial life. The diversity and quantity of microorganisms in soil have also been implied through the analysis of environmental DNA (10). Such analyses highlight the immense variety of microorganisms present in the soil. However, a significant issue is that many of these microorganisms have yet to be successfully cultured (11). The difficulty in growing these microorganisms in artificial media in the lab makes it challenging to fully understand the microbial ecosystems.

Historically, to isolate bacteria from crude samples obtained from environments like soil, the predominant approach has been to culture them on nutrient-rich agar media to promote the growth of individual colonies. However, this method often leads to fast-growing bacterial colonies dominating the culture plates by overshadowing slower-growing ones. This can result in an apparent reduction in colony diversity, limiting the variety of microbes that can be selected for further study. There has been prior research, such as a study by Taha et al. in 2007, which demonstrated that by culturing samples on Soil Compost Agar at 44°C for 7 days, it is possible to efficiently isolate colonies of fast-growing actinobacteria while avoiding the “masking” effect where they overshadow other microbial species (12). It is known that the abundant nutrients in culture media can be toxic to certain types of microorganisms, inhibiting their growth (13). To counteract this, a common strategy is to use diluted media components, which has been shown to improve the cultivation efficiency of environmental bacteria (14). Additionally, the inclusion of supernatants from specific bacterial cultures is known to increase the diversity of bacterial species that develop on agar plates. This technique is utilized to obtain a greater variety of bacterial species from environmental samples (15).

Despite these innovative approaches, the isolation and cultivation of unculturable microbes from the environment have proven to be challenging, with traditional microbial cultivation methods facing significant hurdles. There is a need for the development of alternative media to traditional nutrient-rich media to uncover different aspects of microbial ecology that have not been observed before. In this study, we first created agar media containing extracts from leaf mold to attempt the cultivation of previously uncultured bacteria from soil samples. Furthermore, we performed 16S rRNA gene sequencing analysis on colonies that appeared on the leaf mold media and those that appeared on YME media, to compare the diversity of bacterial strains grown on each type of media.

## Materials and Methods

### Soil Sample Collection

The soil samples used in this study were collected from a park in Tokyo, Japan (Musashino Central Park, Tokyo, Japan, latitude 35° 43’ 7.3776” N, longitude 139° 33’ 30.348” E).

### YME agar plates

The YME agar medium used in this study was prepared as follows: 4g of Yeast extract (Gibco 211929), 10g of Malt extract (Gibco 218630), 4g of D-glucose (Fujifilm 195-01663), and 15g of Agar (Nacalai 01028-85) were added to 1L of reverse osmosis (RO) water. This mixture was then autoclaved at 121°C for 15 minutes. After autoclaving, the medium was dispensed into Petri dishes to solidify, and these were used as the YME agar medium for microbial cultivation.

### Leaf mold medium preparation

leaf mold (product name: leaf mold with Bark, 45 L, from a local supplier, Viva Home Hachioji, headquartered in Saitama Prefecture) was used to prepare the medium. 500 g of the leaf mold was mixed with 2 L of Milli-Q water in a beaker, autoclaved, and then left to stand at room temperature overnight. The liquid extracted by heating was centrifuged, and the supernatant was used as the leaf mold extract. The leaf mold extract was diluted 10-fold with pure water, and 1.5% agar was added. After autoclaving at 121°C for 15 minutes, the mixture was dispensed into plastic petri dishes to create the leaf mold agar plates (Fig. 1).

**Figure 1.**
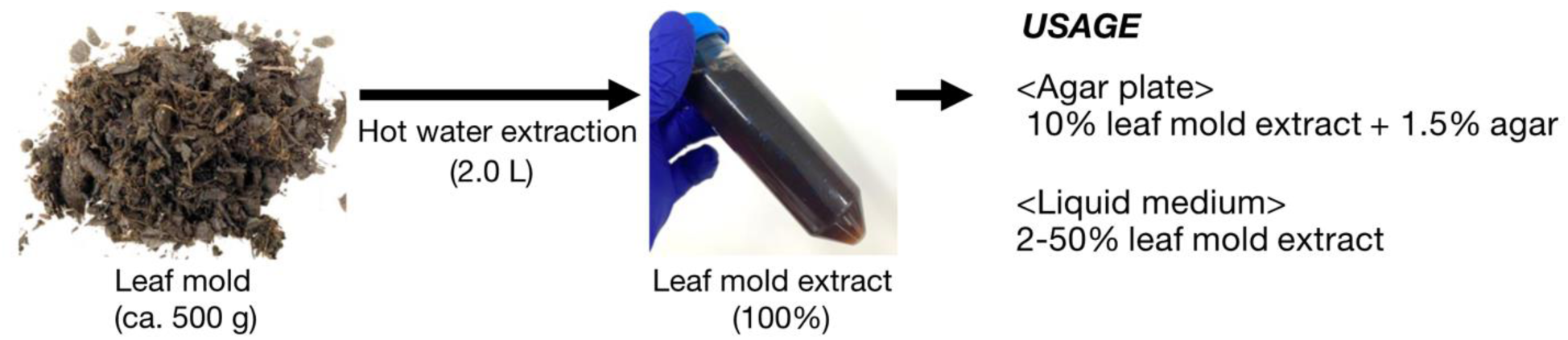
Preparation of Leaf Mold Extract. Commonly used leaf mold (left) was added to Milli-Q water and subjected to hot water extraction by autoclaving to obtain the leaf mold extract (center). This extract was diluted tenfold with Milli-Q water and supplemented with 1.5% agar to be used as leaf mold agar medium. Additionally, the extract was used as a liquid medium in this study.

### Bacterial cultivation using agar media

A small spatula-full of the collected soil sample was suspended in 1 mL of physiological saline, then diluted 10 and 100 times. 100 μL of each dilution was spread onto YME agar and leaf mold agar media, and then incubated at 30 °C for 3 days. From the colonies that grew on the YME and leaf mold media, 68 and 76 colonies were respectively picked. Each was re-streaked onto YME and leaf mold media to isolate single colonies and re-incubated at 30 °C for 2 more days.

### Amplification of DNA Sequences by Colony Direct PCR

Colony direct PCR was performed using colonies isolated on each medium as templates. Premix EX Taq™ Hot Start Version (TaKaRa Bio) was used for the DNA polymerase in the PCR. The primers used for DNA amplification were 16S_8F primer (5’-AGAGTTTGATCCTGGCTCAG- 3’) and 16S_1492R primer (5’-GGTTACCTTGTTACGACTT-3’). The PCR reaction was carried out under conditions modified from the standard protocol provided by the manufacturer: 94°C for 10 min, followed by 30 cycles of 98°C for 10 sec, T°C for 30 sec (T = 45, 50, 55, 60, or 65, depending on sample), and 72°C for 90 sec. Eventually, PCR products were obtained from 59 strains grown on YME media and 57 strains grown on leaf mold agar media.

### 16S rRNA Sequence Analysis

The PCR products obtained were treated and purified with ExoSAP IT (Thermo Fischer Scientific). Briefly, using the purified DNA as a template, DNA sequence analysis was conducted using the 16S_8F primer. In this study, sequence information of 16S rRNA was obtained from 57 strains grown on YME media and 51 strains grown on leaf mold agar media. The approximately 800 bp long DNA sequences obtained were used to identify the genera of the bacteria (please see Table 1 and 2 for actual sequence length for each isolate) grown on the media by conducting a BLAST search on the NCBI rRNA database.

**Table 1.** List of Bacteria Grown on YME Agar Medium.

| No. | Most closely related species | Genus | Read length (bp) | Identity (%) | Identical base pair |
| --- | --- | --- | --- | --- | --- |
| 1 | *Bacillus amyloliquefaciens* | *Bacillus* | 690 | 100 | 690/690 |
| 2 | *Bacillus amyloliquefaciens* | *Bacillus* | 681 | 100 | 681/681 |
| 3 | *Bacillus haynesii* | *Bacillus* | 760 | 99.73 | 758/760 |
| 4 | *Bacillus haynesii* | *Bacillus* | 853 | 99.77 | 851/853 |
| 5 | *Bacillus licheniformis* | *Bacillus* | 863 | 99.65 | 860/863 |
| 6 | *Bacillus licheniformis* | *Bacillus* | 863 | 99.77 | 861/863 |
| 7 | *Bacillus licheniformis* | *Bacillus* | 817 | 100 | 817/817 |
| 8 | *Bacillus mobilis* | *Bacillus* | 842 | 99.88 | 841/842 |
| 9 | *Bacillus mobilis* | *Bacillus* | 797 | 100 | 797/797 |
| 10 | *Bacillus mobilis* | *Bacillus* | 876 | 100 | 876/876 |
| 11 | *Bacillus mobilis* | *Bacillus* | 825 | 99.88 | 824/825 |
| 12 | *Bacillus proteolyticus* | *Bacillus* | 915 | 99.89 | 914/915 |
| 13 | *Bacillus proteolyticus* | *Bacillus* | 940 | 100 | 940/940 |
| 14 | *Bacillus sonorensis* | *Bacillus* | 863 | 99.88 | 862/863 |
| 15 | *Bacillus subtilis* | *Bacillus* | 756 | 99.87 | 755/756 |
| 16 | *Bacillus subtilis* | *Bacillus* | 909 | 100 | 909/909 |
| 17 | *Bacillus subtilis* | *Bacillus* | 901 | 100 | 901/901 |
| 18 | *Bacillus subtilis* | *Bacillus* | 864 | 100 | 864/864 |
| 19 | *Bacillus subtilis* | *Bacillus* | 859 | 100 | 859/859 |
| 20 | *Bacillus subtilis* | *Bacillus* | 899 | 99.89 | 898/899 |
| 21 | *Bacillus subtilis* | *Bacillus* | 884 | 99.89 | 883/884 |
| 22 | *Bacillus tropicus* | *Bacillus* | 854 | 99.77 | 852/854 |
| 23 | *Bacillus zanthoxyli* | *Bacillus* | 898 | 100 (*1) | 898/898 |
| 24 | *Bacillus zanthoxyli* | *Bacillus* | 650 | 100 (*1) | 650/650 |
| 25 | *Bacillus zanthoxyli* | *Bacillus* | 886 | 100 (*1) | 886/886 |
| 26 | *Bacillus zanthoxyli* | *Bacillus* | 908 | 100 (*1) | 908/908 |
| 27 | *Bacillus zanthoxyli* | *Bacillus* | 859 | 100 (*1) | 859/859 |
| 28 | *Bacillus zanthoxyli* | *Bacillus* | 880 | 100 (*1) | 880/880 |
| 29 | *Bacillus zanthoxyli* | *Bacillus* | 882 | 100 (*1) | 882/882 |
| 30 | *Bacillus zanthoxyli* | *Bacillus* | 877 | 100 (*1) | 877/877 |
| 31 | *Bacillus zanthoxyli* | *Bacillus* | 844 | 100 (*1) | 844/844 |
| 32 | *Bacillus zanthoxyli* | *Bacillus* | 872 | 100 (*1) | 872/872 |
| 33 | *Bacillus zanthoxyli* | *Bacillus* | 866 | 99.88 (*2) | 865/866 |
| 34 | *Bacillus zhangzhouensis* | *Bacillus* | 828 | 99.88 | 827/828 |
| 35 | *Bacillus zhangzhouensis* | *Bacillus* | 846 | 100 | 846/846 |
| 36 | *Caballeronia insecticola* | *Caballeronia* | 833 | 100 | 833/833 |
| 37 | *Neobacillus drenthensis* | *Neobacillus* | 841 | 99.64 (*3) | 840/843 |
| 38 | *Paenibacillus favisporus* | *Paenibacillus* | 863 | 99.88 | 862/863 |
| 39 | *Priestia aryabhattai* | *Priestia* | 844 | 100 | 844/844 |
| 40 | *Priestia aryabhattai* | *Priestia* | 697 | 100 | 697/697 |
| 41 | *Priestia aryabhattai* | *Priestia* | 872 | 99.89 (*4) | 871/872 |
| 42 | *Priestia megaterium* | *Priestia* | 873 | 99.89 | 872/873 |
| 43 | *Priestia megaterium* | *Priestia* | 880 | 99.77 | 859/861 |
| 44 | *Priestia megaterium* | *Priestia* | 908 | 100 | 908/908 |
| 45 | *Priestia megaterium* | *Priestia* | 843 | 100 | 843/843 |
| 46 | *Priestia megaterium* | *Priestia* | 883 | 100 | 883/883 |
| 47 | *Rhizobium tibeticum* | *Rhizobium* | 920 | 99.34 | 914/920 |
| 48 | *Streptomyces broussonetiae* | *Streptomyces* | 894 | 99.33 | 888/894 |
| 49 | *Streptomyces caeruleatus* | *Streptomyces* | 901 | 99.89 | 900/901 |
| 50 | *Streptomyces caeruleatus* | *Streptomyces* | 877 | 99.89 | 876/877 |
| 51 | *Streptomyces canus* | *Streptomyces* | 925 | 99.68 | 922/925 |
| 52 | *Streptomyces hachijoensis* | *Streptomyces* | 877 | 99.89 | 876/877 |
| 53 | *Streptomyces hachijoensis* | *Streptomyces* | 874 | 99.77 | 872/874 |
| 54 | *Streptomyces rubiginosohelvolus* | *Streptomyces* | 829 | 100 | 829/829 |
| 55 | *Streptomyces showdoensis* | *Streptomyces* | 861 | 99.07 | 853/861 |
| 56 | *Streptomyces venezuelae* | *Streptomyces* | 878 | 100 | 878/878 |
| 57 | *Streptomyces virginiae* | *Streptomyces* | 894 | 100 | 894/894 |

**Table 2.**
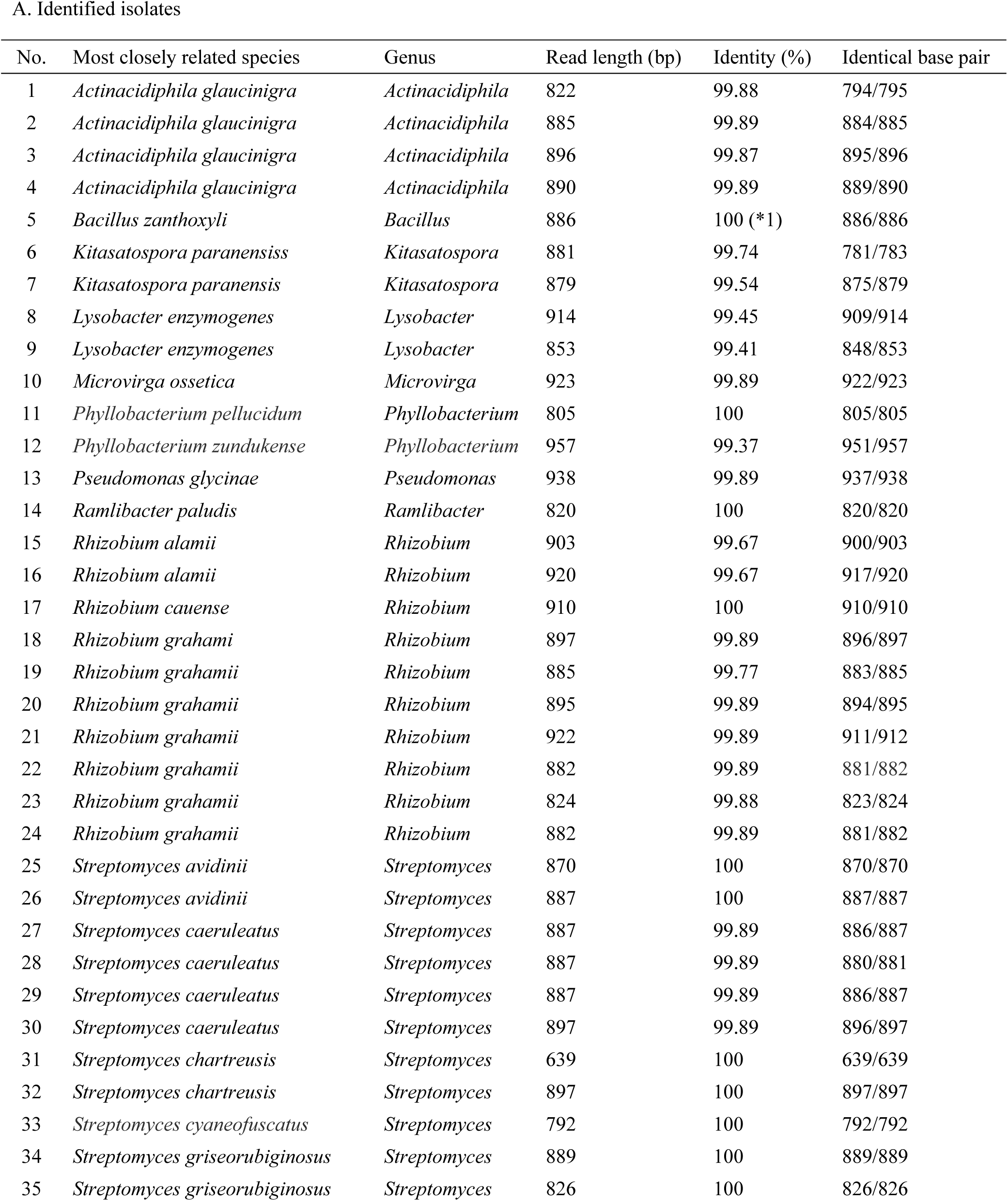
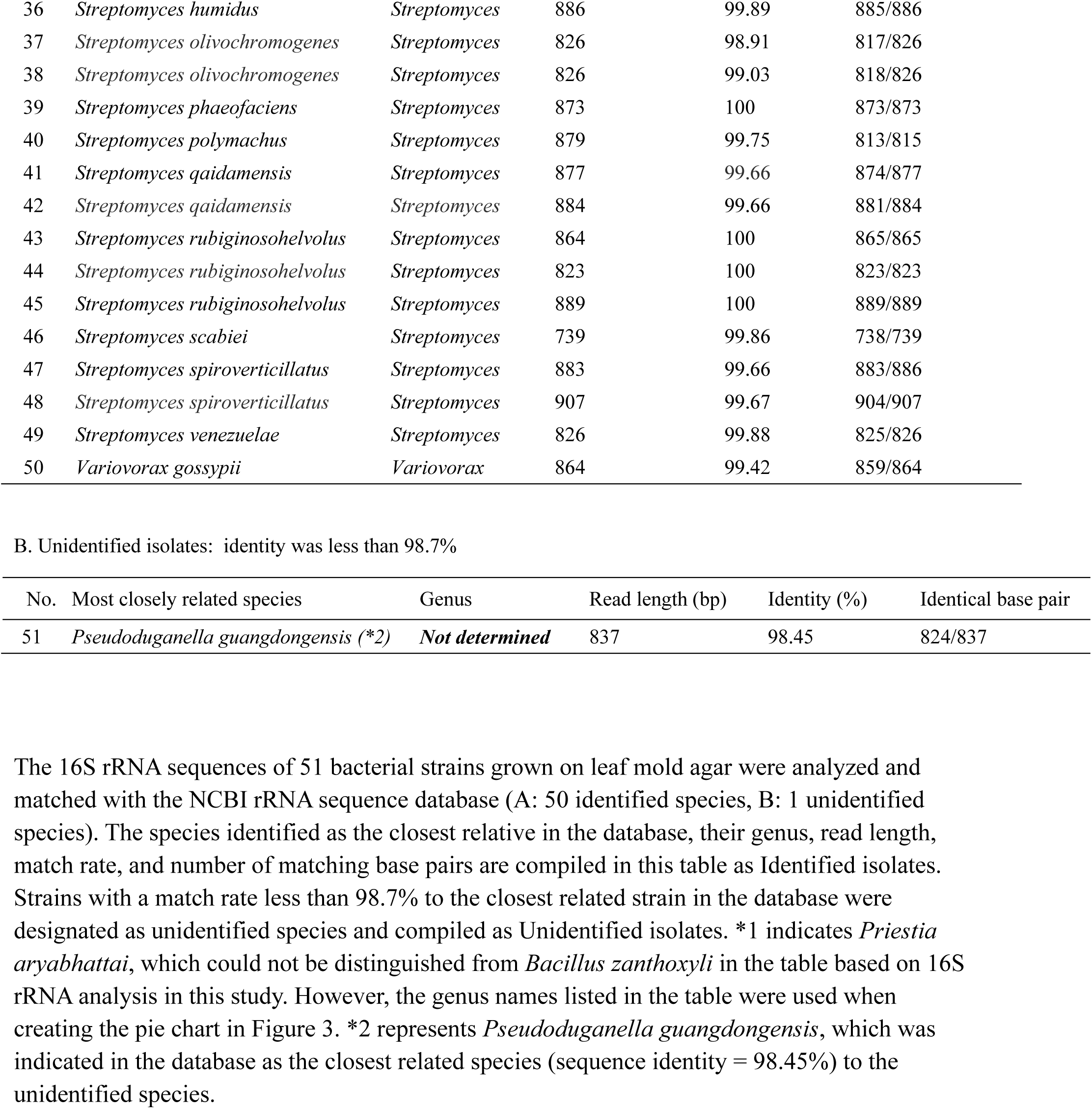
List of Bacteria Grown on Leaf Mold Agar Medium.

### Bacterial Liquid Culture

For the liquid culture of microorganisms, the following media were prepared and used.

*YME (Yeast Malt Extract):* This medium was composed of 4g Yeast extract, 10g Malt extract, and 4g D-glucose dissolved in 1L of reverse osmosis (RO) water. The solution was then autoclaved at 121°C for 15 minutes. *BHI (Brain Heart Infusion):* This medium was prepared by dissolving 37g of Brain Heart Infusion powder (BD, 237500) in 1L of RO water, followed by autoclaving at 121°C for 15 minutes. *LB10 (Luria-Bertani medium):* This medium included 10g Trypton (Gibco 211705), 5g Yeast extract (Gibco 211929), and 10g Sodium chloride (Fujifilm 195-01663) dissolved in 1L of RO water. The solution was autoclaved at 121°C for 15 minutes. *TSB (Tryptic Soy Broth):* This medium was prepared by dissolving 30g of Tryptic Soy Broth powder (BD 211825) in 1L of RO water and autoclaving at 121°C for 15 minutes. *Leaf mold medium:* Additionally, to investigate the effect on the growth of bacteria cultivated on leaf mold agar media, 10% leaf mold extract was prepared (leaf mold exrract : Milli-Q water = 1:9).

## Results

### Differences in Colonies Grown on Media

Soil samples collected from the environment were spread on YME and leaf mold agar media and incubated at 30°C for 3 days, after which the appearance of colonies on each medium was compared (Fig. 2). As a result, on the YME medium, some of the grown colonies were large and spread widely, covering the agar surface. On the other hand, on the leaf mold medium, relatively small colonies were scattered, indicating a significant difference in colony isolation between the two media. These results suggest that using leaf mold agar medium for bacterial culture can yield bacteria that are difficult to obtain with general-purpose nutrient agar media.

**Figure 2.**
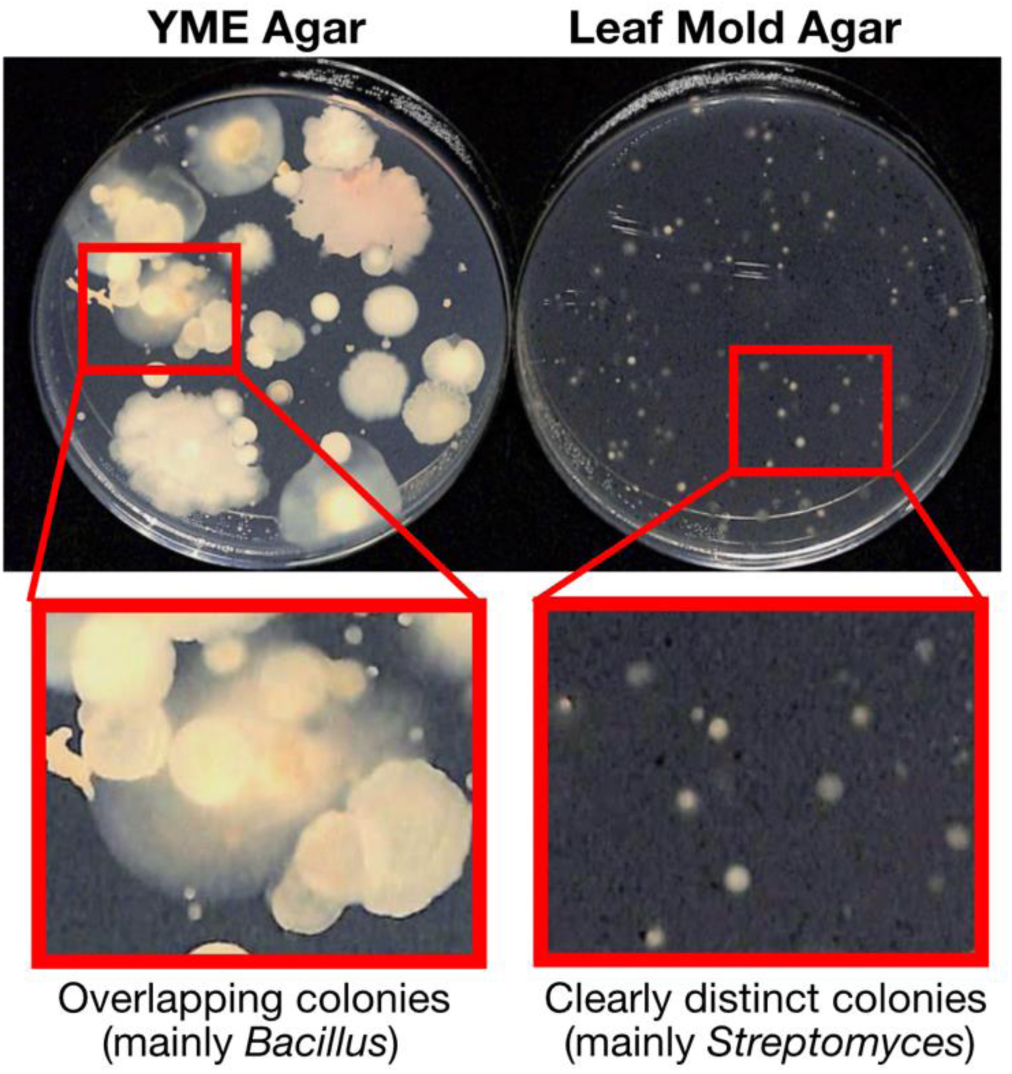
Comparison of Colonies on YME Agar and Leaf Mold Agar Media. A comparison was made of the colony appearance on YME agar and leaf mold agar media after spreading a suspension of soil samples and cultivating at 30°C for 3 days. On YME agar, rapidly growing colonies spread out, overlapping colonies of different bacterial species. In contrast, on leaf mold agar, overlap between colonies was minimized, facilitating isolated culturing.

### 16S rRNA Sequence Analysis of Colonies on Media

Further investigation was conducted to determine if using leaf mold agar medium allows for the isolation of microbes not accessible on YME medium. 16S rRNA sequences of 57 strains grown on YME medium and 51 strains grown on leaf mold agar medium were sequenced and identified at the genus level by matching with database information. The results showed that bacteria isolated from YME medium were predominantly related to the *Bacillus* genus, including *Bacillus*, *Priestia*, *Paenibacillus*, and *Neobacillus*, with actinomyces of the *Streptomyces* genus comprising about 18% of the isolates (Fig. 3 and Table 1). On the other hand, leaf mold agar medium yielded a higher proportion of *Streptomyces* (49%) and rhizobia of the *Rhizobium* genus (20%), along with actinomyces from *Actinacidiphila* and *Kitasatospora* genera, and bacteria from the *Lysobacter* genus (Fig. 3, Table 2). Some bacteria isolated from the leaf mold agar medium showed a similarity of less than 98.7% to the closest related sequence, indicating that species-level identification was not possible based on the 16S rRNA sequences. These results suggest that leaf mold agar medium cultivates a different microbial community from that on YME medium when applied with soil samples.

**Figure 3.**
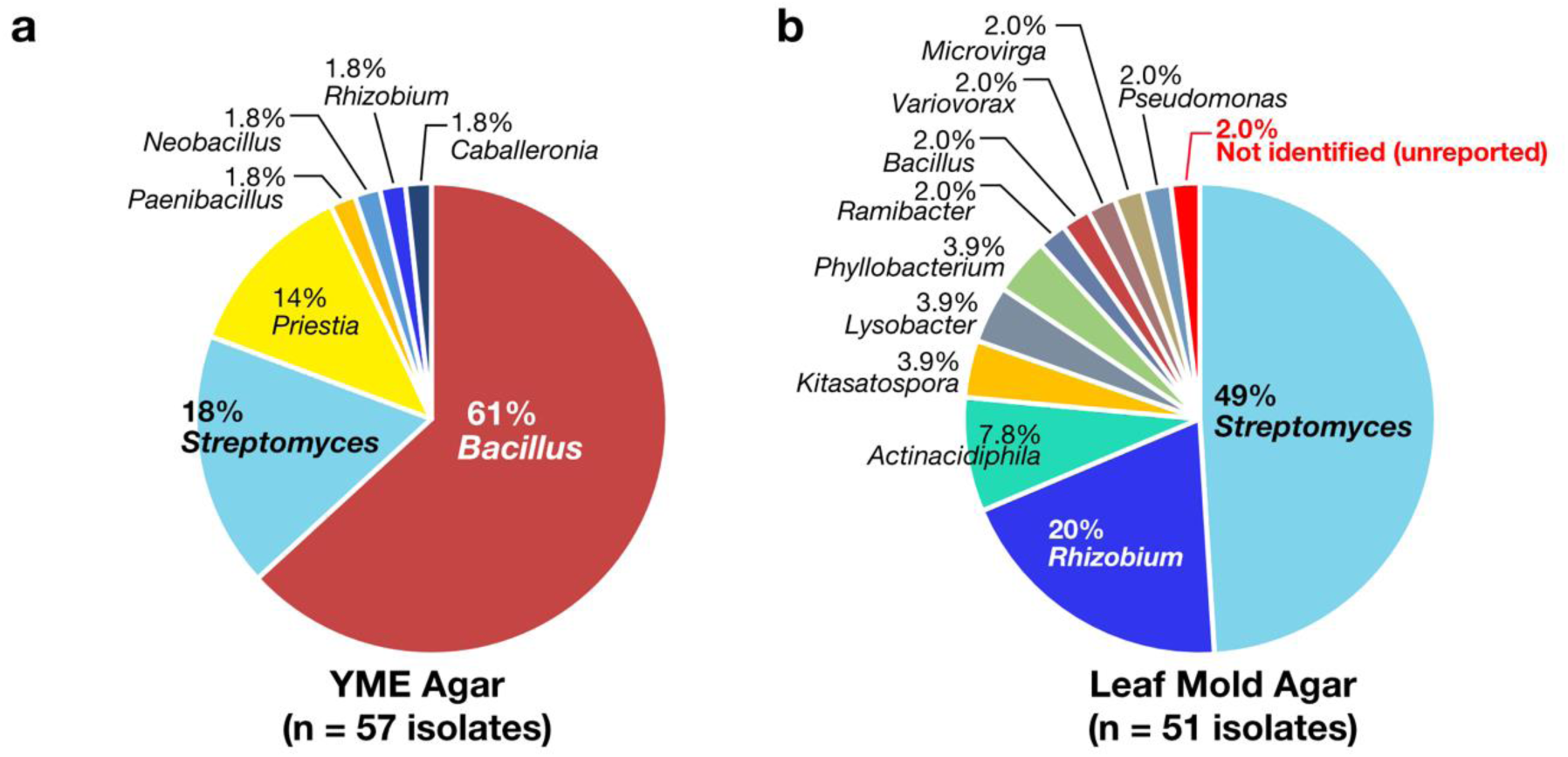
Comparison of Microbial Diversity on YME Agar and Leaf Mold Agar Media. The diversity of microbes grown on (a) YME agar and (b) leaf mold agar was compared by DNA sequencing analysis of the 16S rRNA sequences of the colonies and identifying the bacteria at the genus level using an NCBI database. For detailed information of each isolate, see Tables 1 and 2.

### Liquid Culture of Unidentified Strains Obtained

Colonies of unidentified strain obtained from leaf mold agar medium were inoculated into general nutrient media YME, BHI, LB10, and TSB liquid media to investigate their growth. The results showed no growth in any of the four media (Fig.4). Next, 10% leaf mold extract was prepared to attempt culturing the unidentified strains. The results showed no growth in YME, BHI, LB10, and TSB media with added leaf mold extract, but growth was observed in 10% leaf mold extract medium (Fig.4). These results suggest that adding leaf mold extract enables the cultivation of difficult-to-culture soil bacteria that cannot be grown in general media.

**Figure 4.**
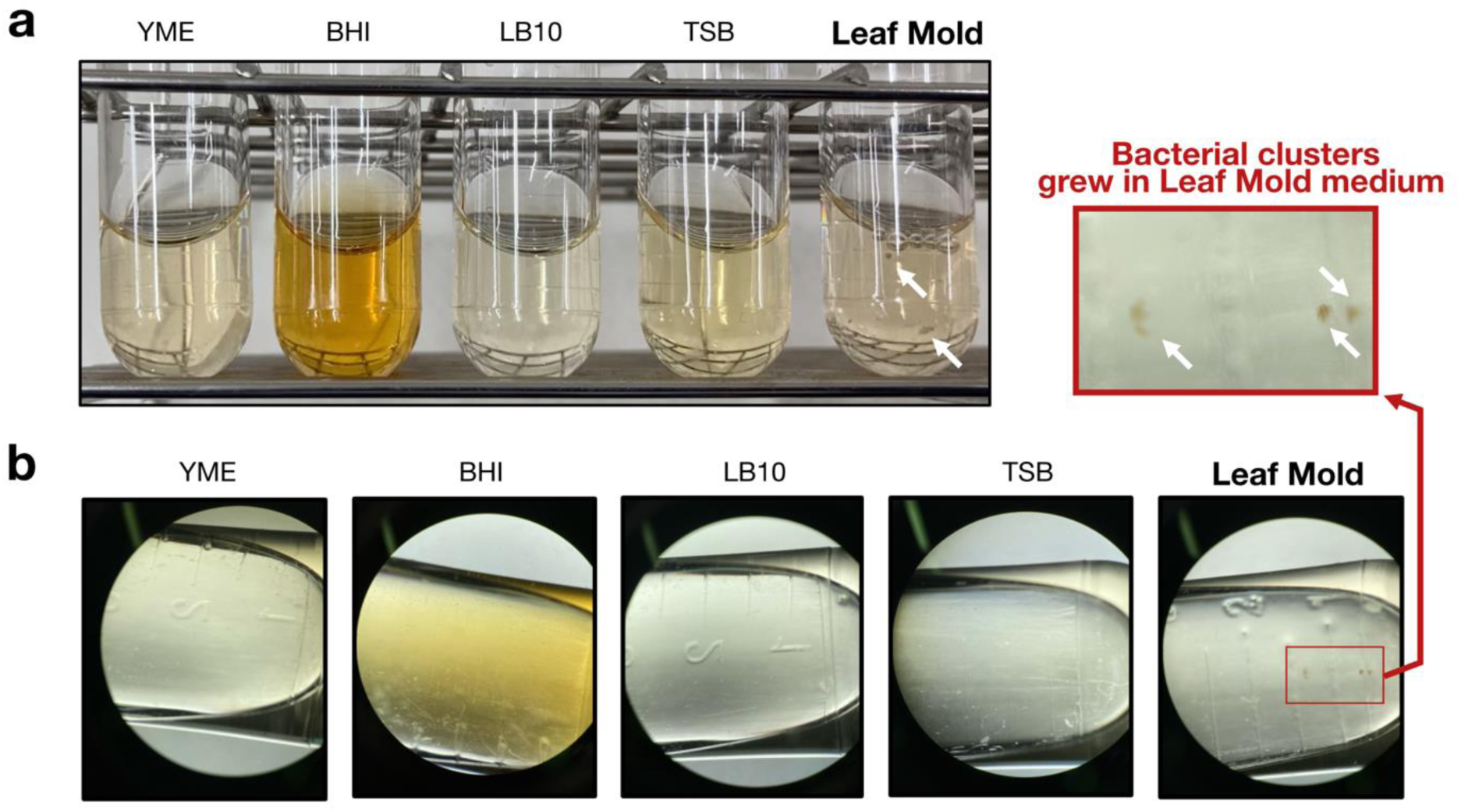
Growth of Isolated Soil Bacteria on Various Nutrient Media. Colonies of an unidentified species grown on leaf mold media were inoculated onto nutrient media such as YME, BHI, LB10, and TSB, or 10% leaf mold extract, and then cultured at 30°C for 3 days. (a) The growth of bacteria after cultivation in test tubes. (b) An observation under a stereomicroscope. Clumps of grown bacteria can be confirmed at the sections indicated by arrows.

## Discussion

In this study, we cultivated bacteria from soil samples using agar media supplemented with leaf mold extract and analyzed their 16S rRNA sequences. We discovered that colonies of bacteria different from those growing on the general-purpose nutrient medium YME can be obtained (Fig. 2 and 3, and Tables 1 and 2). It was also suggested that bacteria that could not be cultured on conventional nutrient media could be cultured by leaf mold extract (Fig. 4). In our laboratory, we are developing experimental systems using invertebrates such as silkworms for applications in drug discovery screening (16–26). However, merely accelerating the assay systems is not sufficient; it is only by constructing a library with high diversity that we can truly enhance the output, that is, exploratory drug discovery research. From this perspective of industrial application, the impact of our research is significant.

Environmental samples have been analyzed through metagenomic and metabolomic analyses to understand the types, quantities, and patterns of secondary metabolites of microbes present (27). However, to understand and actually purify useful components produced by microbes, it is essential to culture them in large quantities in liquid media. This study revealed that the use of leaf mold extract facillitates the cultivation of soil bacteria (Fig. 4). The leaf mold extract contains a set of components needed by soil microbes, and it is expected that culturing, which was difficult with conventional nutrient media alone, becomes possible with the combination of leaf mold extract. In this study, the final concentrations of leaf mold extract added to agar and liquid media were set at 10%, but the optimal concentration, optimal cultivation temperature, cultivation time, shaking speed, and other detailed cultivation conditions for soil microbes remain to be verified in the future study.

Various innovations have been made in the cultivation and examination of microbes from environmental samples. Generally used nutrient media are considered to be excessively nutritious for certain microbes, showing toxicity (13). Therefore, a method to dilute nutrient media by 10 or 100 times for cultivation has been developed (14). Also, there are microbes that depend on secondary metabolites from other biological species not present in artificial media. It has been reported that adding the supernatant of microbial cultures to nutrient media is effective for culturing such microbes (15). Moreover, *in situ* cultivation methods that culture microbes not culturable in artificial media by mimicking real environments are gaining attention (28). This method has reported increased cultivation efficiency in actual environmental samples by returning samples inoculated in cultivation chambers equipped with membranes of impermeable pore size to the environment (29). However, *in situ* cultivation has issues such as the need for specialized devices and difficulty controlling cultivation conditions in natural environments. Despite improvements in cultivation methods, culturing of hard-to-culture microbes has not yet been achieved for many microbes.

The microbial community revealed by the analysis of colonies grown on leaf mold medium was diverse, centered around the *Streptomyces* and *Rhizobium* genera (Fig. 3, Tables 1 and 2). This suggests access to a different pattern of microbial communities than those obtained using conventional media. *Streptomyces* and *Lysobacter* genera, known to produce useful substances like antibiotics (*Streptomyces*: streptomycin and cephamycin C (2, 30); *Lysobacter*: lysocin E (7, 8)), might be accessible through culturing samples collected from various locations using leaf mold media. On the other hand, bacteria obtained from YME agar media were mostly *Bacillus* bacteria. *Bacillus* bacteria hardly grew on leaf mold agar. *Bacillus* bacteria are known to form spores (31). On YME agar media, conditions were suitable for the germination of *Bacillus* spores in soil samples, making *Bacillus* bacteria predominant through cultivation on YME media. On leaf mold agar media, the nutrients necessary for spore germination might not be sufficient, leading to less growth of *Bacillus* compared to YME agar media. Moreover, compounds secreted by dominant microbes on leaf mold agar media, such as *Streptomyces*, might inhibit the growth and spore germination of *Bacillus*. Furthermore, previous studies reported that adding humic acid to media (HV agar) enables selective cultivation of actinomyces from soil (32). Similarly, leaf mold media containing humic components might inhibit the growth of microbes other than actinomyces. Unraveling the mechanisms of these differing growth patterns across media remains a topic for future investigation.

The cultivation of soil samples using leaf mold media in this study yielded a microbial isolate with a 16S rRNA sequence similarity of 98.5%, below the 98.7% criterion for unidentified species (33). Thus, using leaf mold media made it clear that unidentified species can be obtained from soil samples. Unidentified species might possess different genetic resources and patterns of secondary metabolites than conventional microbes. Continued isolation of unidentified species using leaf mold media could be a promising source for the discovery of useful substances, such as pharmaceuticals.

## Declarations

### Author contribution

Conceptualization: AM, MI

Formal analysis: AM, FT

Funding acquisition: AM

Investigation: AM, KM, MM, FT

Project administration: AM, MI

Resources: AM

Supervision: AM

Visualization: AM, FT, MM

Writing – original draft: AM, FT

Writing – review & editing: AM, KM, MI

## Acknowledgment

This work was supported by JSPS KAKENHI (Grant# 22K15461, A.M.), the Research and implementation promotion program through open innovation grants (Grant# JPJ011937, A.M.) from the Project of the Bio-oriented Technology Research Advancement Institution (BRAIN), and Teikyo University Team Resarch Grant (Grant# 22-24, A.M.).

## Competing interest

Authors have no conflict of interest to declare.

## Ethics approval and consent to participate

This study does not include experiments that require ethics approval or consent to participate.

## Consent for publication

All authors checked the final version and agreed to publish this paper.

## Availability of data and material

All data acquired in this study are presented in the manuscript. Additional information and materials are available from the corresponding author upon requests.

